# KLHL9 acts as a KSHV vBcl-2-interacting host factor that supports lytic replication

**DOI:** 10.64898/2026.08.13.744635

**Authors:** Inna Kalt, Sapir Ohev, Anastasia Gelgor Dontsov, Coral Orel Haddad, Itay Koren, Rotem Fuchs, Tzachi Hagai, Ronit Sarid

## Abstract

Kaposi’s sarcoma-associated herpesvirus (KSHV; human herpesvirus 8) is an oncogenic gammaherpesvirus that causes Kaposi’s sarcoma and several lymphoproliferative disorders. The KSHV open reading frame (ORF) 16 encodes viral Bcl-2 (vBcl-2), which inhibits apoptosis and autophagy. vBcl-2 is required for efficient lytic replication and production of infectious progeny; however, this essential role is independent of its established functions regulating cell death. To identify host factors that may contribute to vBcl-2 function, we employed proteomic analysis of HA-vBcl-2 immunoprecipitates from lytically reactivated KSHV-infected cells. This analysis identified the BTB-Kelch protein KLHL9, a substrate-specific adaptor for Cullin 3-RING ligase (CRL3) complex, as a candidate vBcl-2-associated factor. We validated the vBcl-2-KLHL9 association by reciprocal co-immunoprecipitation from infected cells, and by ectopic expression in uninfected cells. During lytic reactivation, KLHL9 redistributed from a predominantly cytoplasmic/perinuclear pattern to a mitochondrial pattern that overlapped with HA-vBcl-2. Alanine-scanning mutagenesis mapped a KLHL9-interaction determinant to the N-terminal region of vBcl-2, and vBcl-2 mutants defective in binding failed to colocalize with KLHL9. Conversely, KLHL9 deletion analysis implicated the C-terminal Kelch-repeat region in efficient association with vBcl-2. AlphaFold modeling supported the interface regions. However, vBcl-2 did not show detectable KLHL9-dependent ubiquitination, suggesting that it is unlikely to be a substrate of a KLHL9-containing complex. Functionally, CRISPR/Cas9-mediated KLHL9 disruption reduced lytic viral protein accumulation and infectious progeny production, while re-expression of sgRNA-resistant KLHL9 partially restored these phenotypes. Together, these findings suggest that vBcl-2 may engage host CRL3 complexes during productive infection, and identify KLHL9 as a vBcl-2-associated host factor that supports efficient KSHV lytic replication.

**Author Summary:** Kaposi’s sarcoma-associated herpesvirus (KSHV) is a cancer-associated virus that alternates between a latent cycle and a productive lytic cycle, during which new infectious virus particles are made. The viral protein vBcl-2 is best known for regulating cell-death and autophagy pathways, but earlier studies showed that it is required for efficient virus production through additional, non-canonical functions. In this study, we identify the cellular protein KLHL9 as a host factor that associates with KSHV vBcl-2. KLHL9 normally functions as a specific adaptor in protein ubiquitination pathways that regulate the turnover, localization, or activity of target proteins through ubiquitination. We show that vBcl-2 and KLHL9 associate with one another, and that KLHL9 redistributes toward the mitochondria in cells expressing vBcl-2. Importantly, loss of KLHL9 impairs the expression of viral lytic proteins and reduces the production of new infectious viruses. This defect resembles the phenotype caused by loss of vBcl-2, suggesting that KSHV employs vBcl-2 to engage KLHL9-dependent host machinery during productive infection. Understanding this interaction may help reveal how herpesviruses reprogram infected cells to create an environment that supports virus production.

## Introduction

Kaposi’s sarcoma-associated herpesvirus (KSHV; human herpesvirus 8) is an oncogenic gammaherpesvirus causally linked to Kaposi’s sarcoma (KS), primary effusion lymphoma (PEL), multicentric Castleman’s disease, and KSHV inflammatory cytokine syndrome [1–7]. Like all herpesviruses, KSHV alternates between latency and lytic replication. During latency, the viral genome persists as a nuclear episome, and viral gene expression is restricted, promoting long-term persistence and immune evasion. In contrast, during the lytic cycle, the replication and transcription activator (RTA) initiates a temporally regulated program of viral gene expression, viral DNA replication, virion assembly, and release of infectious progeny. Lytic infection occurs during primary infection and following reactivation from latency. Although latency predominates in KS lesions and PEL, lytic replication contributes to viral dissemination and elicits inflammatory signals that shape the disease microenvironment. Thus, defining host and viral factors that control the lytic cycle is central to understanding KSHV biology and pathogenesis [8–11].

KSHV open reading frame (ORF) 16 encodes viral Bcl-2 (vBcl-2), a structural and functional homolog of cellular and viral Bcl-2 family proteins that regulate cell survival and cell death pathways [12–17]. Consistent with this homology, vBcl-2 inhibits apoptosis and negatively regulates autophagy [18–21]. However, studies using recombinant bacterial artificial chromosome 16 (BAC16)-derived full-length and vBcl-2-null KSHV genomes demonstrated that vBcl-2 is dispensable for latency, but is required for efficient lytic gene expression, viral DNA replication, and infectious progeny production [22–24]. This lytic function is distinct from the canonical anti-apoptotic and anti-autophagic activities of vBcl-2, as a mutant defective in both cell-death regulatory activities can rescue lytic replication of vBcl-2-null KSHV [23]. Comparative studies further showed that the replication-supporting function is conserved between KSHV and the rhesus rhadinovirus (RRV) vBcl-2 homolog, but is not shared by the more distantly related gammaherpesviruses, murine gammaherpesvirus 68 (MHV68), and herpesvirus saimiri (HVS), the cytomegalovirus mitochondrial inhibitor of apoptosis (vMIA), and cellular Bcl-XL and Bcl-2 proteins [23,24]. Reciprocally, deletion of Bcl-2 from the RRV genome abrogated viral replication, whereas KSHV vBcl-2 rescued RRV replication, indicating that this additional lytic function is conserved between KSHV and RRV [24]. Fine mapping further identified the amino terminus of vBcl-2, particularly glutamic acid 14 (E14), as a critical determinant of its lytic activity. Notably, the E14A mutant retained anti-apoptotic and anti-autophagic functions comparable to those of wild-type vBcl-2 but failed to restore lytic reactivation, highlighting the importance of this residue for KSHV propagation [23].

Subsequent work showed that KSHV vBcl-2 binds the viral tegument protein ORF55 through amino acids 11-20, promotes nuclear localization of both proteins, and facilitates incorporation of ORF55 into the inner tegument. Consistent with the importance of this region, a peptide derived from vBcl-2 residues 11-20 blocked KSHV progeny production without affecting viral gene transcription, protein expression, or viral DNA replication [25]. More recently, vBcl-2 was shown to engage the host nucleoside diphosphate kinase, NM23-H2, to stimulate the GTP loading of the dynamin-related protein 1 (DRP1). This interaction promotes mitochondrial fission, which inhibits aggregation of mitochondrial antiviral signaling protein (MAVS) and impairs the host’s interferon (IFN) response. Critically, the E14A mutant is defective in binding NM23-H2 and fails to induce this mitochondrial remodeling. Collectively, these findings establish vBcl-2 as a multifunctional protein that coordinates cell death, mitochondrial dynamics, immune evasion, and virion architecture to promote productive lytic infection [26].

Cullin-RING ligases (CRLs) are multi-subunit complexes that constitute the largest family of E3-ubiquitin ligases in eukaryotes and regulate diverse cellular processes through proteolytic and non-proteolytic ubiquitin signaling. CRLs are built around a Cullin (CUL) scaffold, whose C-terminal region binds the RING-box protein (RBX1 or RBX2) to recruit the E2 ubiquitin-conjugating enzyme, whereas its N-terminal region engages adaptor and/or substrate-receptor proteins that confer substrate specificity. CRL activity is stimulated by neddylation, the covalent attachment of the ubiquitin-like protein NEDD8 to a conserved C-terminal lysine in the Cullin scaffold, and is further modulated by associated regulatory factors that fine-tune substrate recruitment and ubiquitination [27,28]. Among CRLs, CUL3-containing complexes associate with RBX1 through the CUL3 C terminus and with bric-a-brac/tramtrack/broad complex (BTB/POZ) protein family through the CUL3 N terminus [29–31]. In CUL3 complexes, BTB/POZ proteins function as both adaptors and substrate receptors. A prominent subgroup of these proteins is the BTB-Kelch, or KLHL family, which comprises 42 members. KLHL proteins typically contain an N-terminal BTB domain, a central BACK domain, and a C-terminal Kelch-repeat domain [32]. In addition, a 3-box motif positioned between the BTB and BACK domains cooperates with the C-terminal portion of the BTB domain to engage the N-terminal tail of CUL3 and support CRL3 complex assembly. The Kelch domain, which typically contains five to six Kelch repeats, mediates substrate recognition. Accordingly, deletion of the Kelch repeats does not generally abolish CUL3 binding, whereas deletion of the N-terminal BTB/3-box region disrupts association with CUL3.

Within the KLHL family, the specific functions of KLHL9 remain incompletely defined, but accumulating evidence implicates KLHL9 and its closely related paralog KLHL13 in CUL3-dependent regulation of protein localization, turnover, autophagy-related pathways, and host defense. In humans, a heterozygous KLHL9 missense mutation, p.L95F, located within the conserved BTB domain, has been linked to early-onset autosomal dominant distal myopathy and was shown to reduce interaction with CUL3 [33,34]. KLHL9 and KLHL13 can also function together in a CRL3-KLHL9/KLHL13 complex that monoubiquitylates Aurora B kinase, promoting its relocalization from mitotic chromosomes to spindle microtubules, thereby supporting cytokinesis [35,36]. KLHL9 has been further implicated in proteasomal regulation of the transcription factors C/EBPbeta and C/EBPdelta in mesenchymal glioblastoma [37]. In host-defense pathways, the CRL3-KLHL9/KLHL13 complex interacts with CLEC12A, a receptor involved in sensing danger and pathogen-associated molecular patterns, and contributes to antibacterial autophagy [38]. More recently, *Burkholderia pseudomallei* was shown to hijack the CRL3-KLHL9/KLHL13 complex to promote K63-linked ubiquitination and autophagic turnover of the mitochondrial inner membrane protein IMMT, thereby inducing mitophagy, reducing mitochondrial reactive oxygen species, and enhancing intracellular bacterial survival [39]. Finally, deficiency of the autophagy factor ATG16L1 increases KLHL9 and KLHL13 expression, promoting CRL3-dependent ubiquitination and proteasomal degradation of insulin receptor substrate 1 (IRS1), with consequent impairment of insulin signaling [40].

In the present study, we identify KLHL9 as a cellular factor associated with KSHV vBcl-2, map the domains that mediate this interaction, demonstrate vBcl-2-depedent redistribution of KLHL9 to mitochondria, and show that KLHL9 functions as a proviral host factor that promotes productive KSHV lytic replication.

## Results

### Generation of a recombinant KSHV genome expressing HA-tagged vBcl-2

To identify vBcl-2-associated proteins during the KSHV lytic cycle, we generated a recombinant KSHV genome encoding N-terminally HA-tagged vBcl-2 (BAC16-HA-vBcl-2) using two-step Red recombination (Fig. 1A-C) [22,41,42]. Recombinant virus was reconstituted by transfecting HEK293T cells with wild-type BAC16 or BAC16-HA-vBcl-2 DNA, followed by hygromycin selection and co-cultivation with iSLK cells after treatment with TPA and sodium butyrate to induce lytic reactivation.

**Figure 1.**
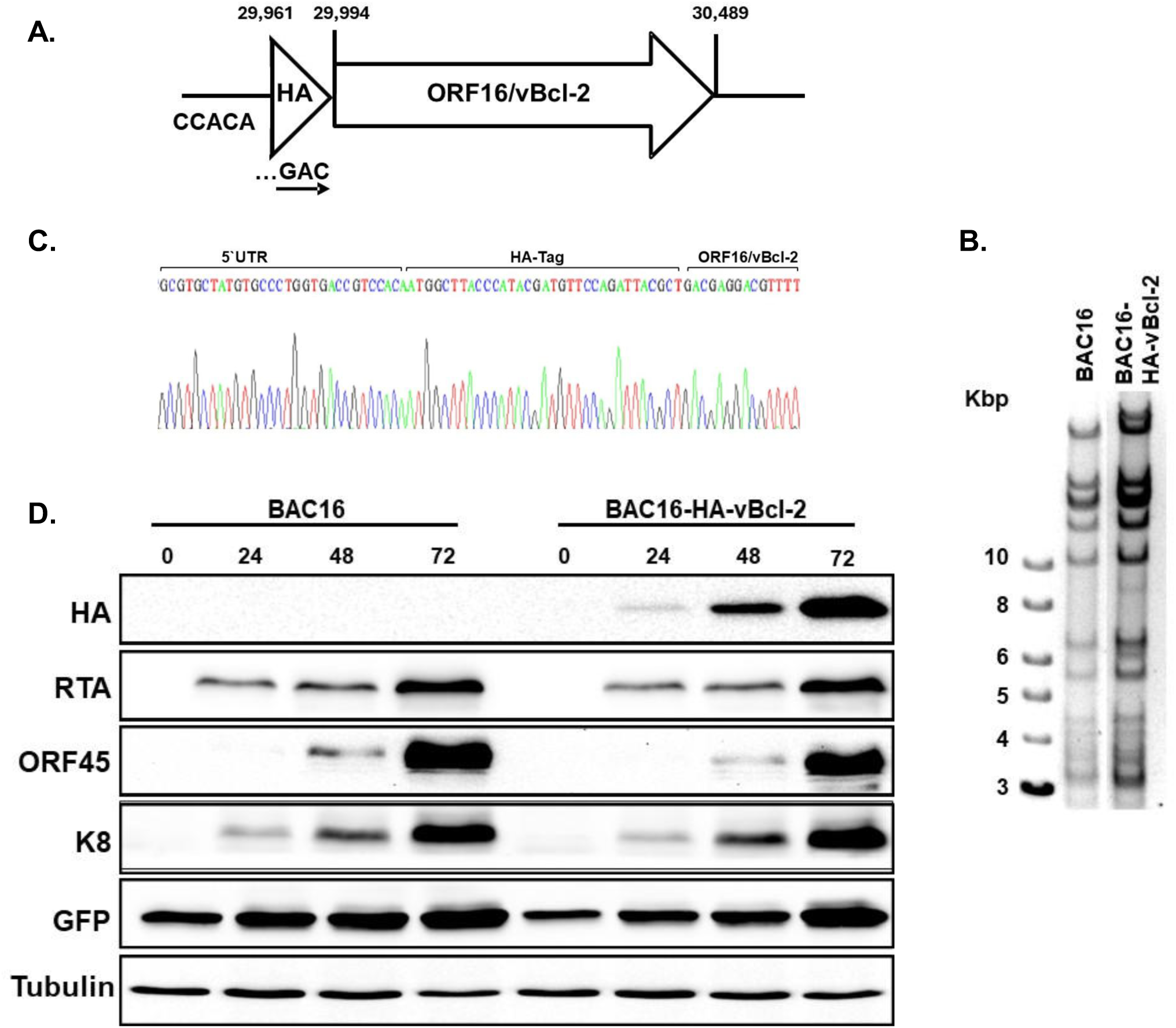
Construction and characterization of recombinant BAC16 expressing N-b terminally HA-tagged vBcl-2 (BAC16-HA-vBcl-2). **(A)** A schematic of the recombinant KSHV BAC16 ORF16/vBcl-2 locus showing insertion of the N-terminal tag (BAC16-HA-vBcl-2). Nucleotide positions refer to the KSHV BAC16 sequence, GenBank accession number GQ994935.1. The positions of the vBcl-2 initiation codon (ATG) and stop codon (29,993 and 30,489) are indicated above the diagram. **(B)** Agarose gel electrophoresis of wild-type BAC16 and BAC16-HA-vBcl-2 DNA digested with BamHI and resolved on a 0.4% ethidium bromide-stained agarose gel. Molecular size markers are shown on the left, with sizes indicated. **(C)** Sequence analysis of the HA insertion site, including the upstream ORF16/vBcl-2 5′ untranslated region, the initiation codon, the HA-tag coding sequence, and the ORF16/vBcl-2 coding region. **(D)** iSLK cells harboring parental BAC16 or BAC16-HA-vBcl-2 were induced into the lytic cycle with sodium butyrate and doxycycline. At 0, 24, 48, and 72 h post induction, whole-cell lysates were prepared and analyzed by immunoblotting with the indicated antibodies against viral proteins and GFP. Tubulin served as loading control. A representative experiment from two independent experiments with similar results is shown.

To assess whether the HA tag detectably affects lytic gene expression, iSLK cells harboring BAC16-HA-vBcl-2 or parental wild-type BAC16 were induced into the lytic cycle, and whole-cell lysates were collected 24, 48, and 72 h postinduction for immunoblot analysis. As shown in Fig. 1D, HA-vBcl-2 was readily detected with an anti-HA antibody following lytic induction, and migrated at an apparent molecular mass of approximately 20 kDa in lysates from BAC16-HA-vBcl-2-infected cells. Re-probing the blot for the viral lytic proteins RTA, ORF45, K8, and ORF65, as well as GFP, revealed comparable expression kinetics and overall levels between BAC16-HA-vBcl-2 and wild-type BAC16-infected cells. These results are in line with previous reports [24,26] and indicate that N-terminal HA tagging does not detectably impair lytic reactivation.

### Proteomic identification and validation of KLHL9 as a vBcl-2-associated protein

To identify proteins that associate with vBcl-2 during lytic virus reactivation, we immunoprecipitated HA-vBcl-2 from BAC16-HA-vBcl-2-infected iSLK cells following induction of the lytic cycle, and analyzed the precipitates by liquid chromatography-tandem mass spectrometry (LC-MS/MS). Induced parental BAC16-infected cells were processed in parallel as a negative control. Across two independent experiments, 50 viral and cellular proteins were enriched at least 5-fold in HA-vBcl-2 immunoprecipitates relative to control samples (S1 Table). Among the highest-scoring candidates was KLHL9, a BTB-Kelch protein and candidate substrate receptor for CRL3 complexes. Notably, a previous genome-wide interaction screen also identified KLHL9 as a candidate vBcl-2-associated protein in cells ectopically expressing vBcl-2, supporting its prioritization for validation [43]. Furthermore, given the reported roles of KLHL9 in mitotic regulation, autophagy and host responses to pathogens, we considered KLHL9 a strong candidate mediator of noncanonical vBcl-2 functions.

We validated the association between vBcl-2 and KLHL9 by co-immunoprecipitation. In HEK293T cells infected with BAC16-HA-vBcl-2 and ectopically expressing KLHL9-Flag during lytic reactivation, KLHL9-Flag co-immunoprecipitated with HA-vBcl-2 following anti-HA immunoprecipitation (Fig. 2A), and reciprocal immunoprecipitation with anti-Flag antibody recovered HA-vBcl-2 in KLHL9-Flag precipitates (Fig. 2B). We further examined whether endogenous KLHL9 associates with vBcl-2 during lytic reactivation. Using an antibody against KLHL9/13, an endogenous KLHL9/13-reactive protein co-immunoprecipitated with HA-vBcl-2 from BAC16-HA-vBcl-2-infected iSLK cells induced into the lytic cycle (Fig. 2C). Finally, in uninfected HEK293T cells, transiently expressed HA-vBcl-2 co-precipitated with KLHL9-Flag as well as with endogenous KLHL9/13-reactive protein, indicating that the interaction does not require additional viral proteins (Fig. 2D). Together, these results identify KLHL9 as a vBcl-2-associated cellular factor, and validate the interaction in both infected cells and uninfected ectopic-expression systems.

**Figure 2.**
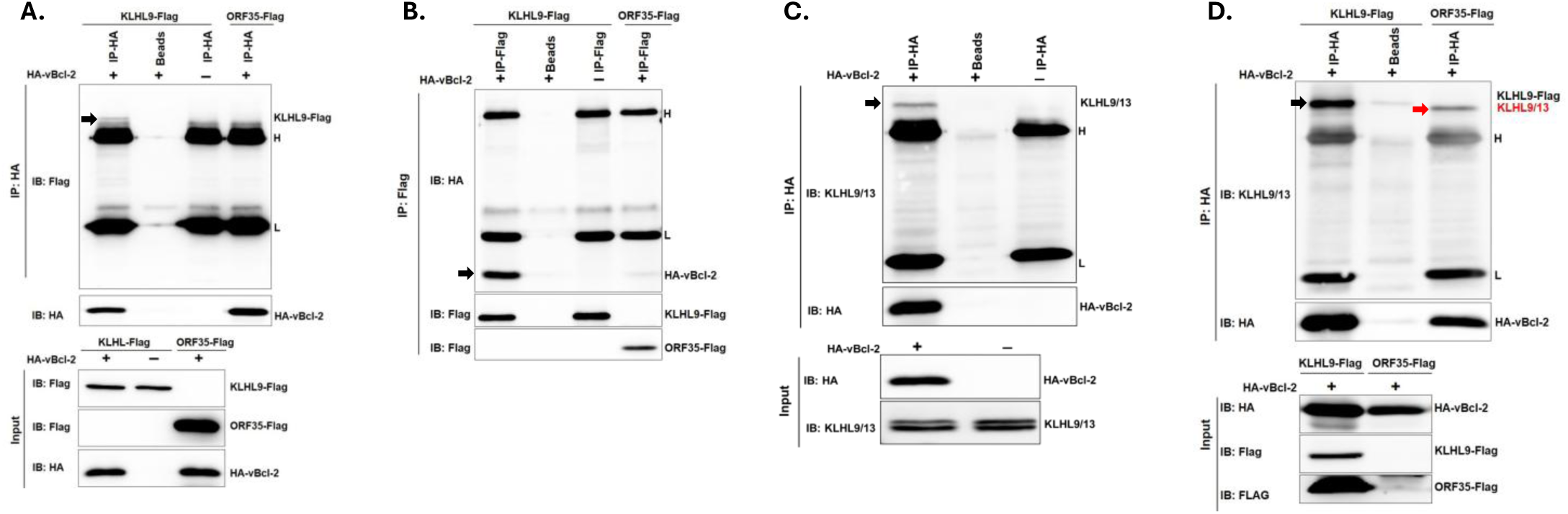
vBcl-2 associates with KLHL9 in infected and uninfected cells. HEK293T cells infected with recombinant BAC16-HA-vBcl-2 (HA-vBcl-2+) or parental BAC16 (HA-vBcl-2−) were transfected with KLHL9-Flag or, as a negative control, ORF35-Flag. After 24 h, lytic reactivation was induced with TPA and sodium butyrate. Cells were harvested 72 h later, and protein extracts were subjected to immunoprecipitation (IP) with anti-HA antibody followed by Flag immunoblotting **(A)** or anti-Flag antibody followed by HA immunoblotting **(B).** Co-immunoprecipitated proteins are indicated by black arrows. Membranes were also probed with anti-HA **(A)** or anti-FLAG **(B)** antibodies to confirm HA-vBcl-2, KLHL9-Flag and ORF35-Flag expression (Input) and immunoprecipitation. **(C)** Co-immunoprecipitation of endogenous KLHL9/13-reactive protein with HA-vBcl-2 during lytic reactivation. BAC16-HA-vBcl-2-infected iSLK cells were induced into the lytic cycle with doxycycline and sodium butyrate for 72 h, and protein extracts were immunoprecipitated with anti-HA antibody. Immunoblotting was then performed with anti-KLHL9/13 antibody to detect co-immunoprecipitated endogenous protein, indicated by a black arrow. The membrane was re-probed with anti-HA antibody to confirm HA-vBcl-2 expression. **(D)** Uninfected HEK293T cells were transiently transfected with expression plasmids encoding HA-vBcl-2 and KLHL9-Flag; ORF35-Flag served as a negative control. After 48 h, cells were harvested and protein extracts were subjected to immunoprecipitation with anti-HA antibody followed by immunoblotting with anti-KLHL9/13 antibody to detect co-immunoprecipitated proteins, indicated by black and red arrows. Membranes were also probed with anti-HA, anti-KLHL9/13, and anti-Flag antibodies to confirm protein expression. Protein extracts incubated with beads alone served as a negative control. L and H indicate antibody light and heavy chains, respectively. Representative blots are shown from three independent experiments with similar results.

### KLHL9 localizes with vBcl-2 during lytic reactivation

KSHV vBcl-2 was previously reported to localize predominantly to mitochondria during lytic reactivation [24,26], whereas KLHL9 displays dynamic intracellular localization depending on cell type and context [35,39,44]. To determine the subcellular localization of KLHL9 during KSHV lytic reactivation, BAC16-HA-vBcl-2-infected iSLK cells were transduced with a lentiviral vector encoding mCherry-KLHL9, and KLHL9 localization was analyzed by immunofluorescence using an anti-mCherry antibody. As shown in Fig. 3, in uninduced cells, mCherry-KLHL9 was predominantly cytoplasmic with pronounced perinuclear accumulation. At 48 h after lytic induction, KLHL9 redistributed to the mitochondria and showed extensive colocalization with HA-vBcl-2, which predominantly colocalized with the mitochondria as well. These observations support the vBcl-2-KLHL9 association in live cells and suggest that lytic reactivation is accompanied by redistribution of KLHL9 to vBcl-2-positive mitochondrial regions.

**Figure 3.**
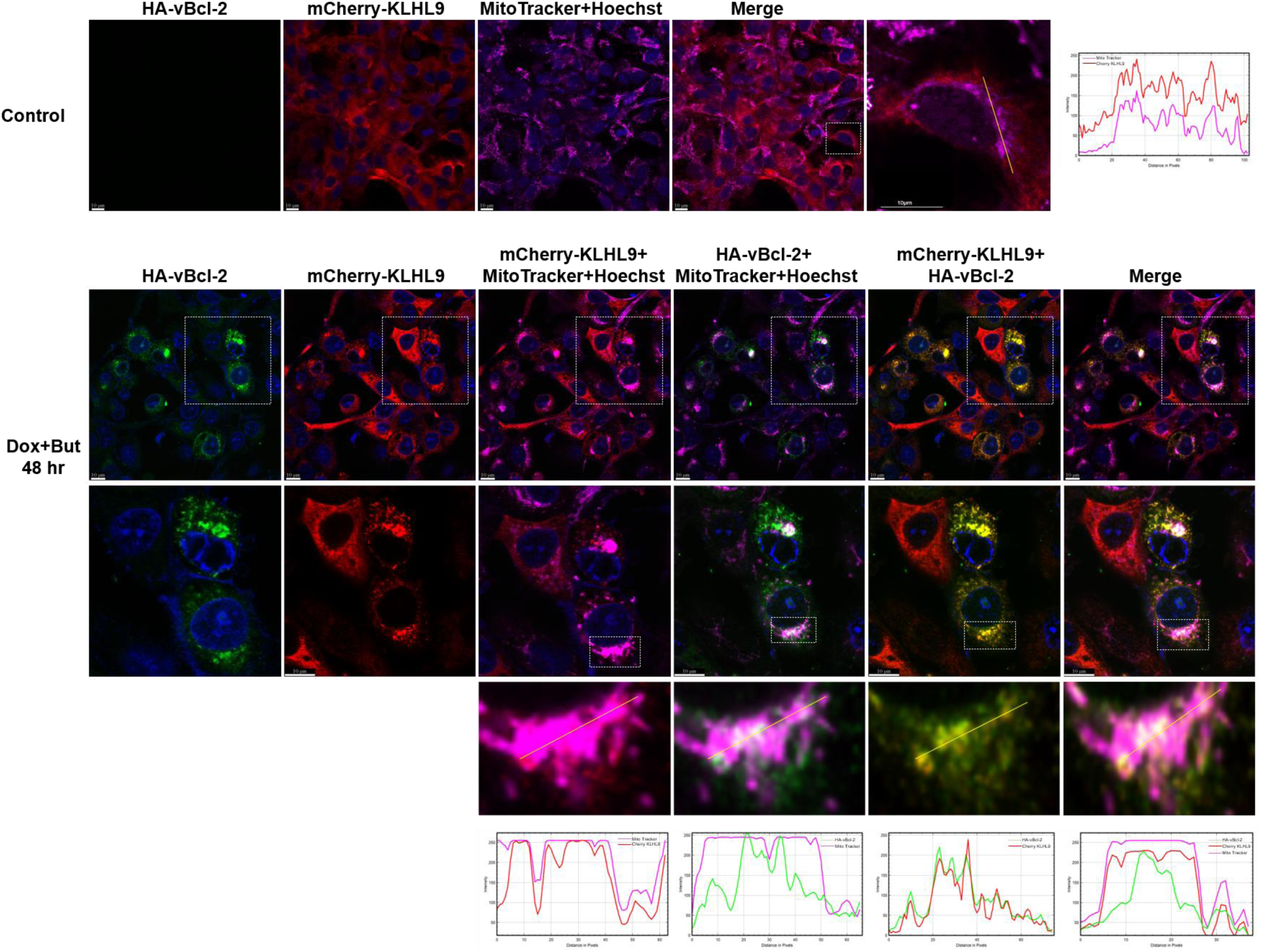
KLHL9 re-localizes with vBcl-2 during lytic reactivation. BAC16-HA-vBcl-2-\ infected iSLK cells expressing mCherry-KLHL9 were left uninduced (control), or induced with doxycycline and sodium butyrate for 48 h. Mitochondria were labeled with MitoTracker Deep Red (magenta), and mCherry-KLHL9 (red) and HA-vBcl-2 (green) were detected by immunofluorescence using anti-mCherry and anti-HA antibodies, respectively, followed by Cy3 and 488 secondary antibodies, respectively. Nuclei were stained with Hoechst. Scale bar, 10 µ. Boxed areas are shown underneath in an enlarged view. The right most upper panel and bottom panels show the fluorescence intensity profiles measured along the line drawn by ImageJ.

### The vBcl-2 N-terminal region is required for KLHL9 association

To map the region of vBcl-2 required for interaction with KLHL9, we used a previously described set of pcDNA-HA-vBcl-2 alanine-scanning mutants spanning amino acids 1-139 [23]. In each mutant, a block of five consecutive residues was substituted with alanines (Fig. 4A). We transiently co-transfected HEK293T cells with plasmids encoding wild-type or mutant HA-vBcl-2 along with mCherry-KLHL9. Cells were harvested 48 h after transfection, and whole-cell lysates were subjected to anti-HA immunoprecipitation followed by immunoblotting with anti-mCherry antibody. As shown in Fig. 4B, mutants A4-A8, corresponding to residues 16-40, failed to interact with KLHL9. In addition, the HA-vBcl-2-E14A mutant, which was previously shown to disrupt vBcl-2 interactions with ORF55 and NM23-H2 [25,26], retained binding to KLHL9. Notably, Liang et al. previously reported that A3-A4 and A6-A8 fail to rescue, or only partially rescue, lytic replication of vBcl-2-null virus, suggesting that these N-terminal regions contribute to the essential lytic function of vBcl-2. Together, these findings map the KLHL9-binding determinant of vBcl-2 to a discrete N-terminal region encompassing residues previously implicated in the noncanonical lytic functions of vBcl-2.

**Figure 4.**
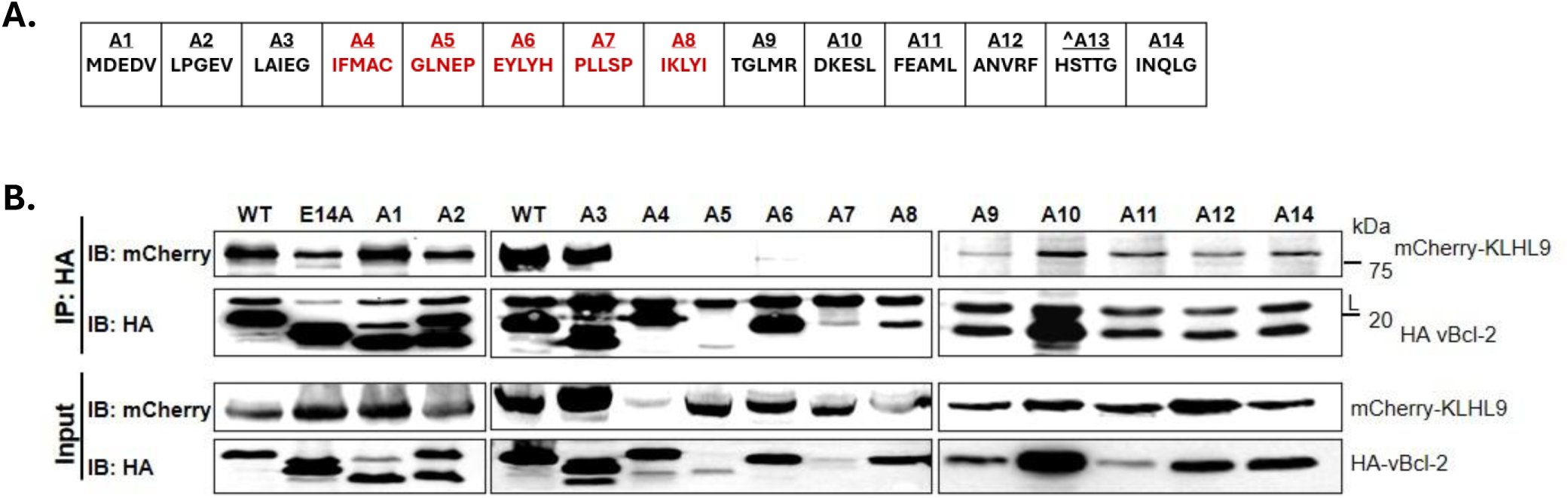
N-terminal residues of vBcl-2 contribute to KLHL9 association. (A) Schematic representation of selected alanine-scanning mutants of HA-vBcl-2 spanning amino acids 1-139 (A1-A14). Each mutant contains substitution of a block of four to five consecutive residues with alanines. **(B)** HEK293T cells were transiently co-transfected with mCherry-KLHL9, and the indicated wild-type or mutant HA-vBcl-2 expression plasmids. Mutant A13 was excluded from the analysis and the HA-vBcl-2 E14A mutant was included for comparison. Cells were harvested 48 h after transfection, and whole-cell lysates were subjected to immunoprecipitation with anti-HA antibody followed by immunoblotting with anti-mCherry antibody. Mutants A4-A8, corresponding to residues 16-40 did not interact with KLHL9.

To further characterize the vBcl-2 mutants defective in KLHL9-binding, we examined their intracellular localization relative to the mitochondria and mCherry-KLHL9. HEK293T cells were transiently transfected with expression plasmids encoding wild-type HA-tagged vBcl-2 or the interaction-defective A4-A8 mutants, either alone or together with mCherry-KLHL9. Wild-type HA-vBcl-2 colocalized with mitochondria, and generated an altered distribution of mitochondria compared with control untransfected cells (Fig. 5A). Wild-type HA-vBcl-2 also showed clear colocalization with mCherry-KLHL9 (Fig. 5B). Among the interaction-defective mutants, mutants A6 and A8 retained mitochondrial localization and the ability to alter the cellular distribution of the mitochondria similar to wild-type vBcl-2, whereas the remaining mutants showed reduced mitochondrial colocalization and did not detectably alter mitochondrial distribution. Consistent with the co-immunoprecipitation results, mutants that failed to interact with KLHL9 were also unable to colocalize with mCherry-KLHL9.

**Figure 5.**
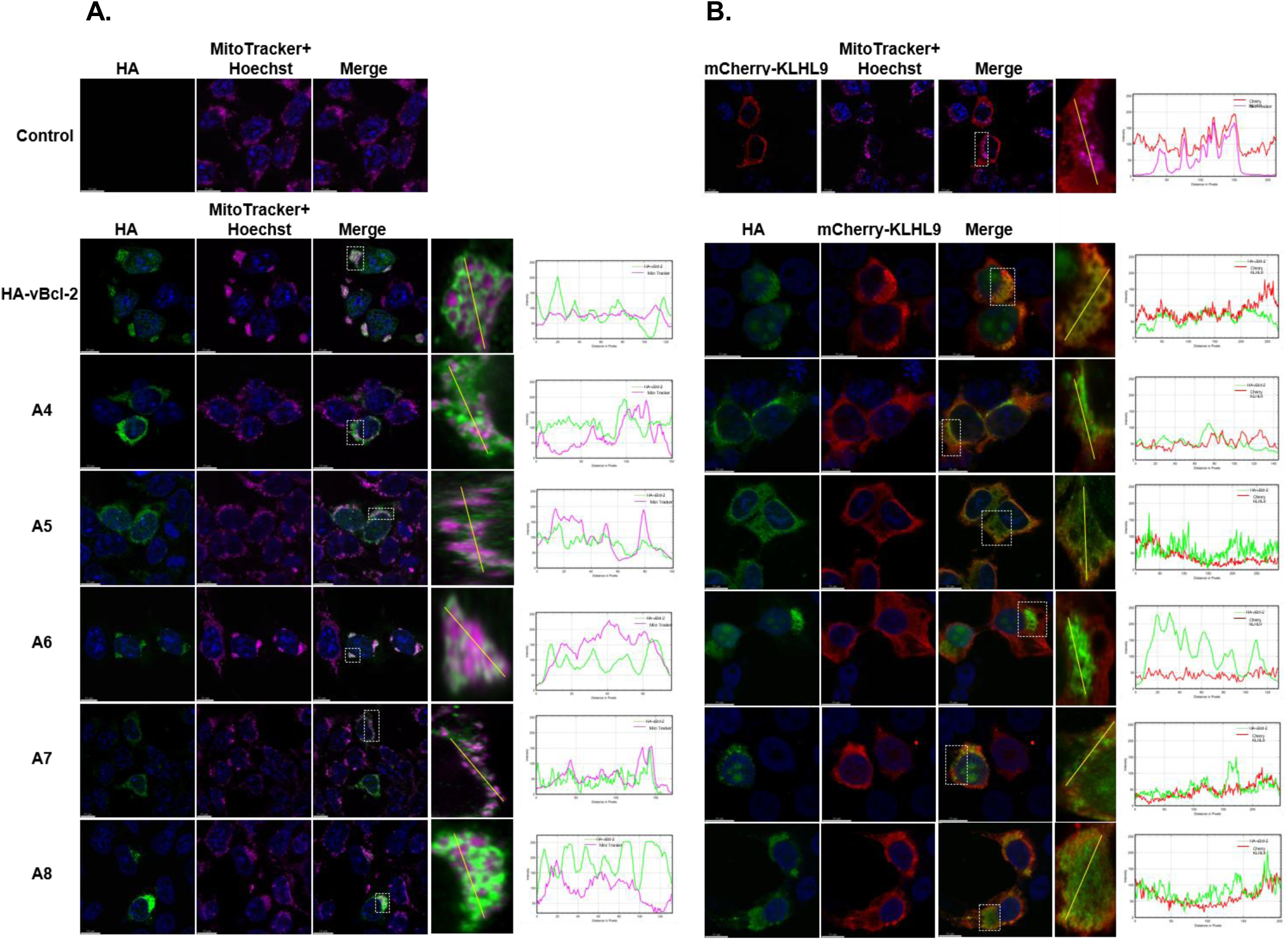
vBcl-2 mutants defective in KLHL9 association fail to colocalize with KLHL9. **(A)** HEK293T cells were transiently transfected with expression plasmids encoding wild-type or mutant HA-vBcl-2 proteins (A4-A8). Control cells were transfected with vector only. Mitochondria were labeled with MitoTracker, and wild-type and mutant HA-vBcl-2 proteins were detected with anti-HA antibody followed by an Alexa Fluor 488-conjugated secondary antibody. Boxed areas are enlarged, with the right most panels showing the fluorescence intensity profiles measured along the line drawn by ImageJ. **(B)** HEK293T cells were transfected with mCherry-KLHL9 alone (control) or co-transfected with mCherry-KLHL9, and expression plasmids encoding wild-type or mutant HA-vBcl-2 (A4-A8). At 48 h post transfection, cells were fixed and analyzed by immunofluorescence. mCherry-KLHL9 was detected with anti-mCherry antibody followed by a Cy3-conjugated rabbit secondary antibody, and HA-vBcl-2 proteins were detected with anti-HA antibody followed by an Alexa Fluor 488-mouse conjugated secondary antibody. Boxed areas are enlarged with the right-most panels showing the fluorescence intensity profiles measured along the line drawn by ImageJ.

Together, these observations indicate that the vBcl-2 N-terminal region contributes to KLHL9 association, and suggest that KLHL9 binding is separated from at least some aspects of vBcl-2 mitochondrial localization and remodeling.

### The KLHL9 Kelch-repeat region is required for efficient vBcl-2 association

BTB-Kelch proteins often use their C-terminal Kelch-repeat domains for substrate or partner recognition [32]. To determine which region of KLHL9 contributes to vBcl-2 association, we generated mCherry-tagged KLHL9 deletion constructs (Fig. 6A) and co-expressed them with HA-vBcl-2 in HEK293T cells. Co-immunoprecipitation analysis showed that constructs containing the Kelch-repeat region associated efficiently with vBcl-2, whereas constructs lacking this region failed to associate with vBcl-2 (Fig. 6B). These results implicate the KLHL9 Kelch-repeat domain in vBcl-2 association. Consistent with our results, AlphaFold2-based structural modeling suggested that the interaction interface involves the Kelch-repeat regions of KLHL9 and the N-terminal region of vBcl-2 (Fig.7). We note that the predicted interface is composed of a relatively large number of residues, including residues within the N-terminal alanine-substitution vBcl-2 mutants that failed to bind KLHL9, as detailed above. The binding residues from KLHL9 include several regions from different loops that are distant in primary sequence but are spatially close, likely forming a dynamic interface with vBcl-2. The complete list of interacting residues is detailed in Supporting S2 Table.

**Figure 6.**
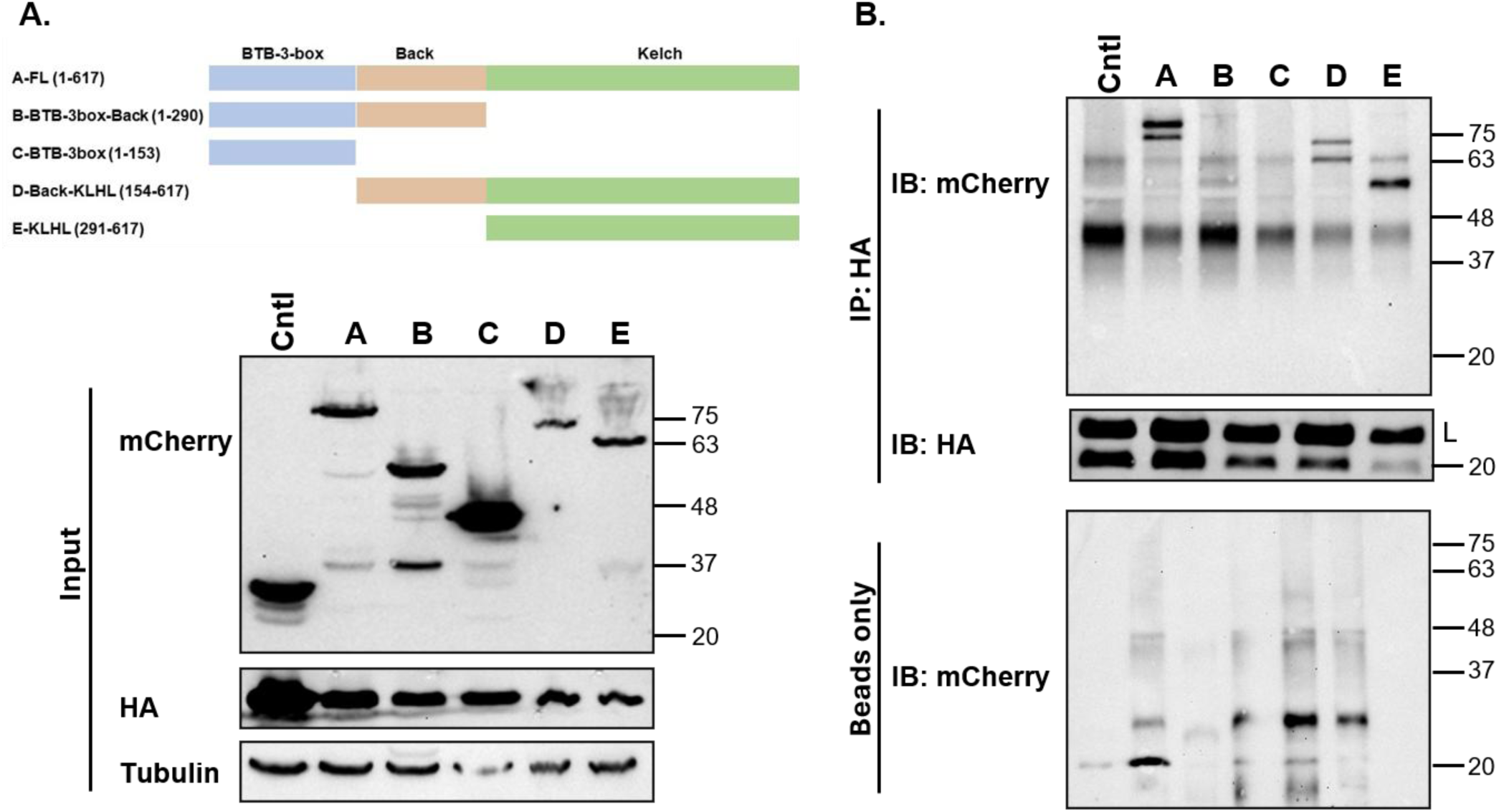
Interaction of vBcl-2 with KLHL9 requires the Kelch-repeat domain. **(A)** Schematic representation of the KLHL9 deletion mutants used for domain-mapping analysis. **(B)** HEK293T cells were co-transfected with HA-vBcl-2, and the indicated mCherry-tagged KLHL9 construct. Expression of the transfected proteins was verified by immunoblotting of whole-cell lysates (Input). Cell lysates were then subjected to immunoprecipitation with anti-HA antibody, and the immunoprecipitates were analyzed by immunoblotting with anti-mCherry and anti-HA antibodies to detect KLHL9 derivatives and vBcl-2, respectively. Lysates from cells co-transfected with mCherry alone (Cntl), as well as samples incubated with agarose beads alone (Beads only), served as negative controls.

**Figure 7.**
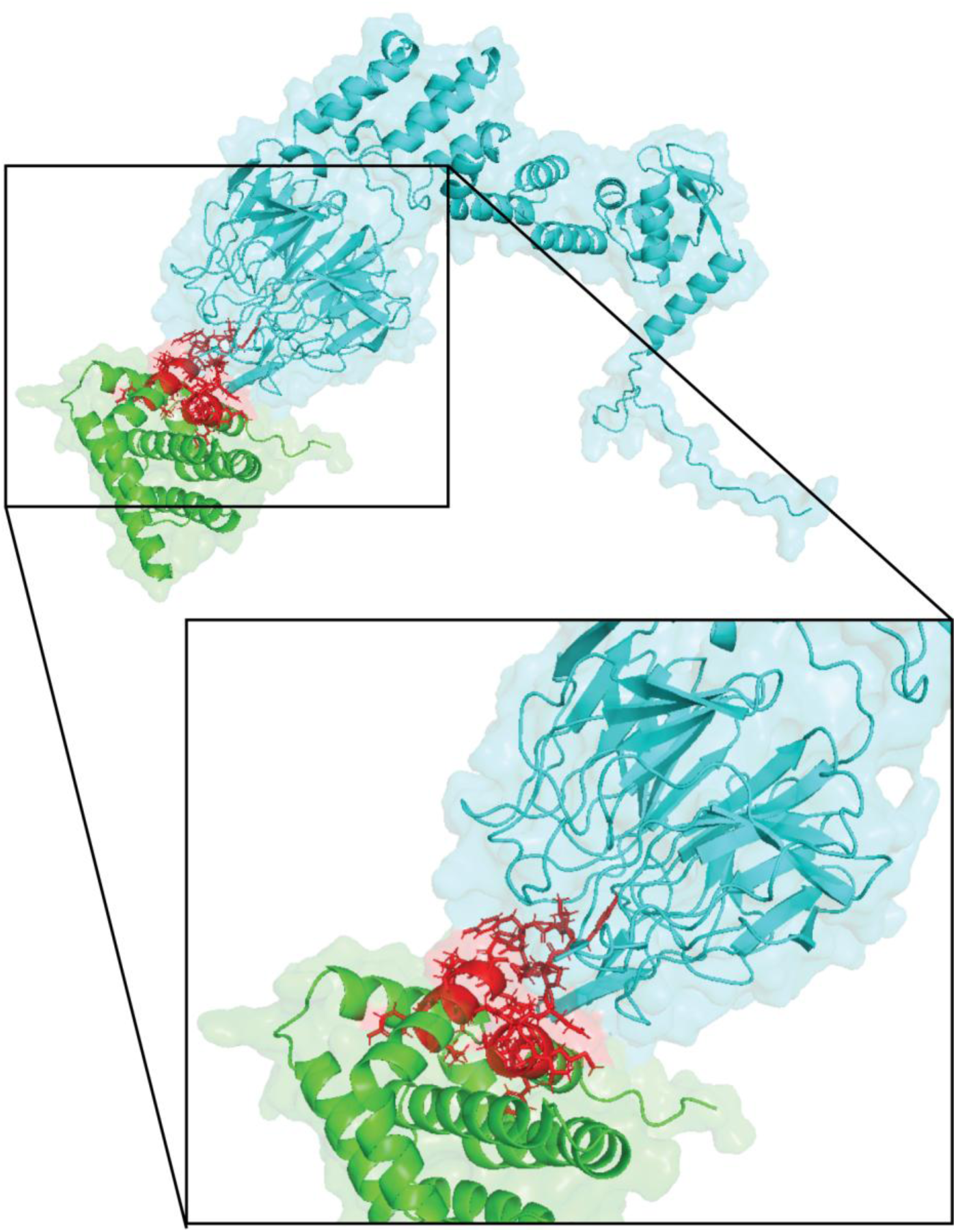
Structural modelling of the predicted protein complex formed between vBcl-2 and KLHL9. An AlphaFold-multimer predicted structure is shown. vBcl-2 and KLHL9 are marked in green and cyan, respectively. Predicted interface residues are shown in red. An enlarged version is shown to the right, focused on the interface region. Both figures were created using PyMol (https://www.pymol.org/). The complete list of interacting residues is detailed in Supporting S2 Table.

### vBcl-2 is not detectably ubiquitinated by KLHL9

Because KLHL9 can function as a substrate adaptor for the CRL3 complexes, we tested whether vBcl-2 is detectably ubiquitinated in a KLHL9-dependent manner. As a non-lysine acceptor control, we generated a vBcl-2 mutant in which all four lysine residues were replaced with arginine (vBcl-2 K/R). HEK293T cells were co-transfected with HA-vBcl-2 or the HA-vBcl-2 K/R variant together with mCherry-KLHL9 and Myc-tagged Ubiquitin, followed by treatment with either bortezomib or MG132 to inhibit proteasomal degradation. Under these conditions, we did not detect evidence of enhanced ubiquitination of vBcl-2 in the presence of KLHL9 (Fig.8). These data argue against a simple model in which vBcl-2 is the primary KLHL9-dependent ubiquitination substrate, but do not exclude ubiquitin-dependent regulation of other viral or cellular substrates.

**Figure 8.**
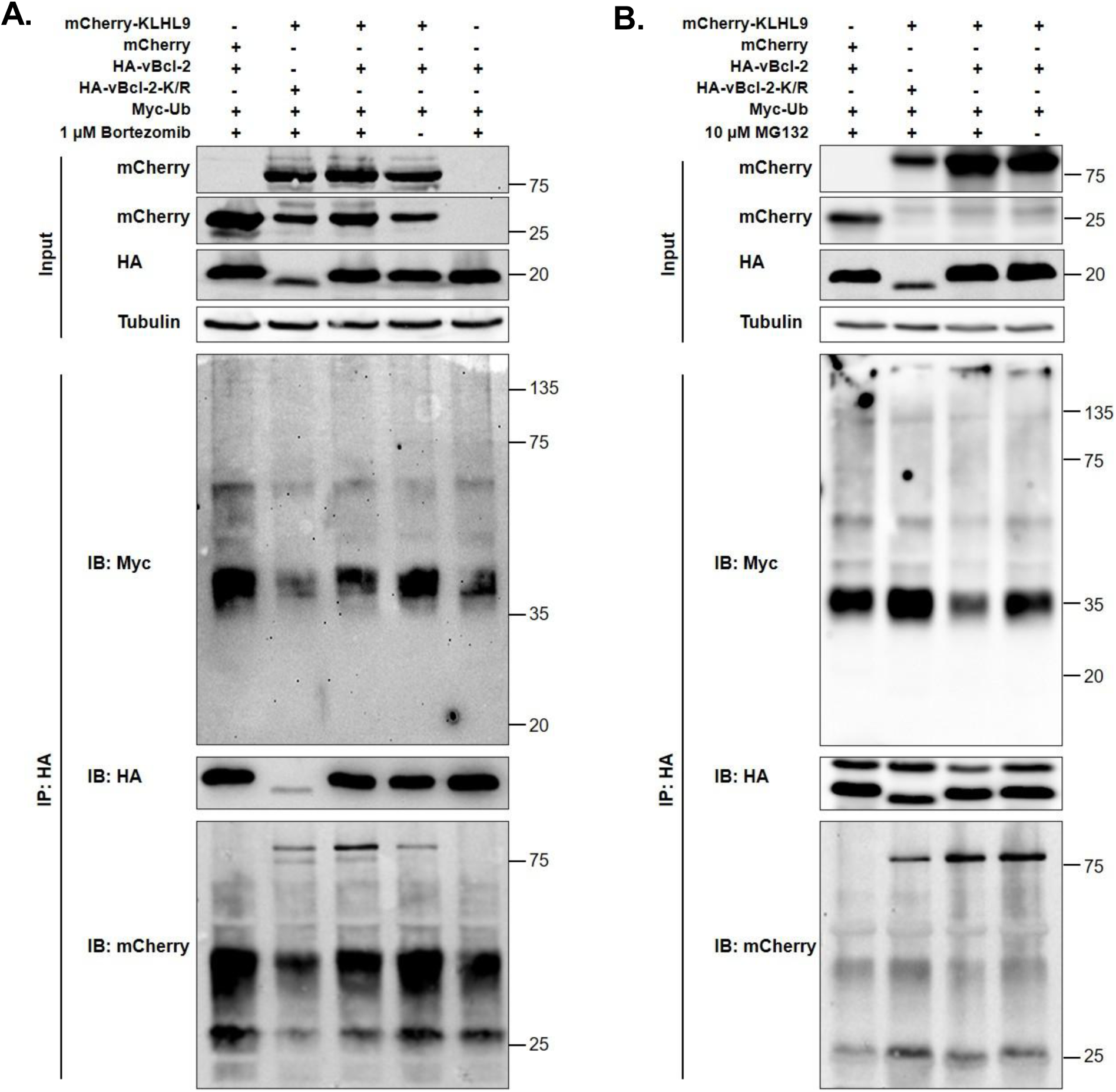
vBcl-2 is not detectably ubiquitinated by KLHL9. HEK293T cells were co-transfected with HA-vBcl-2 or HA-vBcl-2 K/R, in which all four lysine residues were replaced with arginine, together with mCherry-KLHL9 or mCherry vector control and Myc-tagged Ubiquitin. At 48 h post-transfection, cells were treated with 1 µM bortezomib for 6 h **(A)**, or 10 µM MG132 for 4 h **(B)** and then harvested. Whole-cell extracts were prepared in the presence of 70 µM PR619 deubiquitinase inhibitor, and analyzed by immunoblotting with the indicated antibodies (Input). HA-tagged proteins were immunoprecipitated with anti-HA antibody, and immunoprecipitates were analyzed by immunoblotting with the indicated antibodies.

### KLHL9 supports efficient KSHV lytic reactivation and infectious progeny production

Having established that vBcl-2 interacts with KLHL9, we next examined the contribution of KLHL9 to the KSHV infection cycle. To this end, we used CRISPR/Cas9 to target endogenous KLHL9 in BAC16-HA-vBcl-2-infected iSLK cells, using a single-guide RNA (sgRNA) cloned into the LentiCRISPRv2 vector. Successful knockout was confirmed by sequencing, and yielded a frameshift mutation predicted to introduce a premature stop codon. To assess specificity of the knock-out, we performed rescue experiments by re-expressing KLHL9 from a lentiviral vector encoding mCherry and an sgRNA-resistant KLHL9-Flag, carrying synonymous substitutions within the sgRNA target sequence. Control cells were transduced with the corresponding mCherry-expressing empty vector.

We then assessed the effect of KLHL9 deficiency on KSHV lytic reactivation by examining the expression of selected viral lytic proteins and quantifying infectious progeny production. As shown in Fig. 9A, B, KLHL9 knockout resulted in reduced lytic viral protein expression and a markedly decreased production of infectious progeny. Similar results were obtained using an independent knockout iSLK cell population. Interestingly, this phenotype is reminiscent of that previously observed in cells infected with vBcl-2-null KSHV [22–24]. Re-expression of sgRNA-resistant KLHL9 partially restored lytic viral protein expression and infectious virus production, confirming that KLHL9 contributes functionally to the KSHV lytic program (Fig. 9A,B). Consistent with these results, attenuation of lytic replication was also observed in KLHL9/KLHL13 double-knockout HEK293T cells (Fig. 9C). Collectively, these findings identify KLHL9 as a proviral host factor that promotes efficient KSHV lytic reactivation, and support a model in which KLHL9 cooperates with vBcl-2 during the lytic cycle.

**Figure 9.**
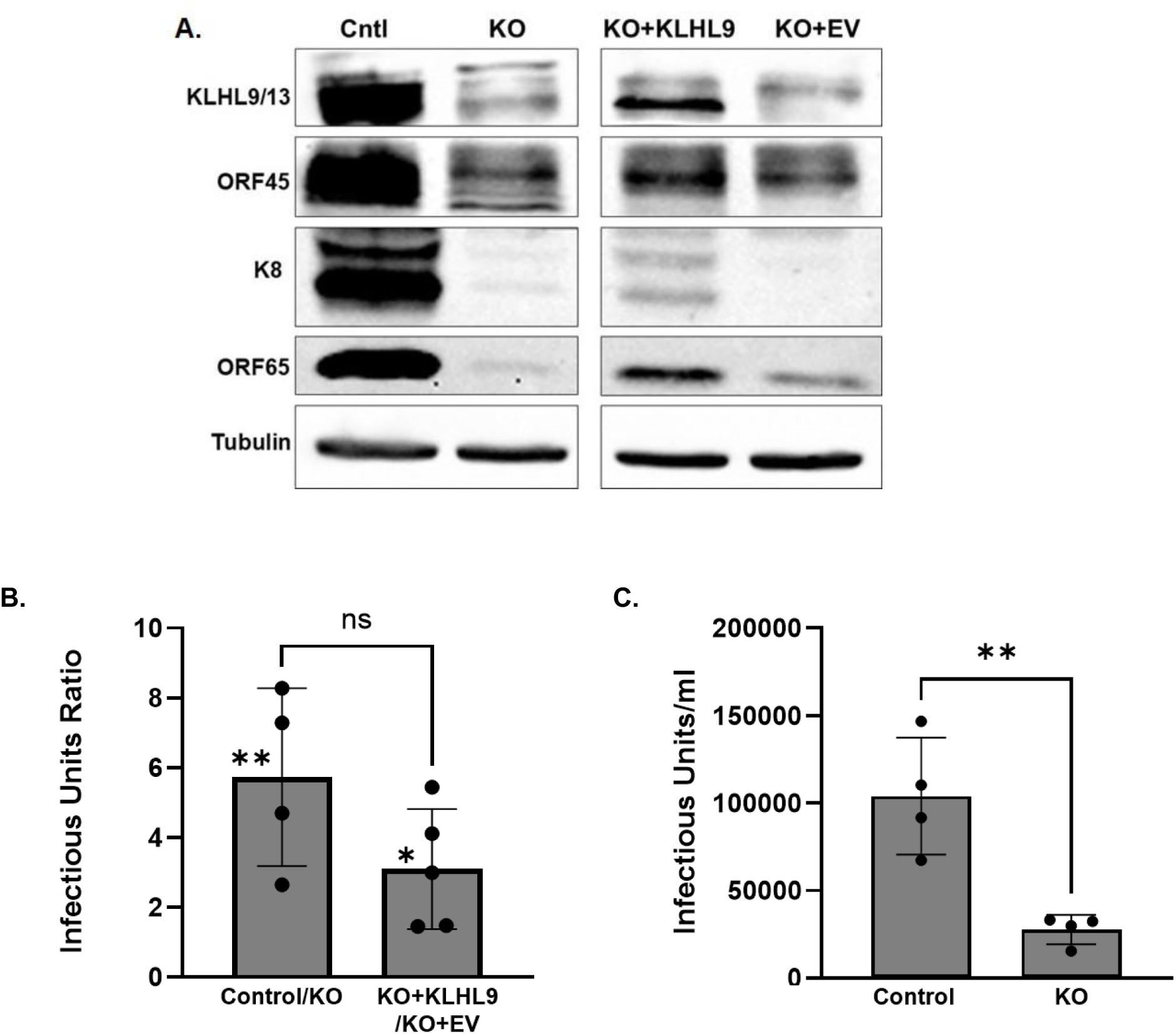
KLHL9 supports efficient KSHV lytic reactivation and infectious progeny production. **(A)** BAC16-HA-vBcl-2-infected iSLK cells were transduced with recombinant lentiviruses encoding either an sgRNA targeting KLHL9 (KO) or a non-targeting control (Cntl). KLHL9-knockout iSLK cells were complemented with a lentiviral vector expressing sgRNA-resistant KLHL9-Flag (KO+KLHL9) or with the corresponding control vector (KO+EV). Cells were induced into the lytic cycle with doxycycline and sodium butyrate for 72 h, and viral lytic protein expression was analyzed by immunoblotting. **(B)** Culture supernatants were collected, and infectious virus production was quantified by flow-cytometric analysis of GFP-positive cells following infection of naive SLK cells. Results are presented as the ratio of infectious virus yield in Cntl versus KO cells and in KO+KLHL9 versus KO+EV. Points represent independent biological replicates; bars indicate mean +/-SD/SEM. **(C)** Control and KLHL9/KLHL13 double-knockout HEK293T cells infected with BAC16-HA-vBcl-2 KSHV were induced into the lytic cycle with TPA and sodium butyrate for 72 h, and infectious virions released into the culture medium from the HEK293T cultures were quantified. Points represent independent biological replicates; bars indicate mean +/− SD/SEM.

## Discussion

In this study, we identify KLHL9 as a host factor associated with KSHV vBcl-2 and provide evidence that KLHL9 supports efficient KSHV lytic replication. KLHL9 emerged from a proteomic analysis of HA-vBcl-2 immunoprecipitates obtained during lytic reactivation, and the interaction was validated by reciprocal co-immunoprecipitation in infected cells, by co-precipitation with endogenous protein, and in ectopic expression assays performed in the absence of other viral proteins. Of note, global mapping of herpesvirus-host protein complex study identified KLHL9 as a potential vBcl-2 partner using proteomic analysis of affinity purified Strep-tagged vBcl-2 in HEK293T cells [43]. In parallel, KLHL9 re-localized from a mainly cytoplasmic/perinuclear distribution to mitochondria during lytic induction and co-localized with vBcl-2. Furthermore, this mitochondrial localization of KLHL9 was dependent on its interaction with vBcl-2. Functionally, disruption of KLHL9 markedly reduced lytic viral protein accumulation and infectious progeny production, while re-expression of sgRNA-resistant KLHL9 partially restored these phenotypes. The fact that KLHL9 deficiency phenocopied the previously described vBcl-2-null defect is notable, because it links a newly identified cellular interactor to the same stage of the viral infection cycle already known to depend on vBcl-2. These observations support a model in which KLHL9 acts as a proviral host factor that is functionally coupled to vBcl-2 during lytic infection.

Our mapping experiments further refine the relationship between KLHL9 association and the noncanonical lytic role of vBcl-2. The KLHL9-binding determinant mapped to the N-terminal region of vBcl-2, with alanine-scanning mutants spanning residues 16-40 showing undetectable interaction and loss of colocalization with KLHL9. Importantly, these same regions overlap sequences previously implicated in the essential lytic function of vBcl-2 [23]. At the same time, the E14A mutant retained KLHL9 binding, despite its previously reported functional defects including its inability to rescue lytic replication [23], its defect in ORF55 interaction and virion assembly [25], and in NM23-H2/DRP1-linked mitochondrial remodeling [26]. This separation-of-function suggests that KLHL9 binding alone is not sufficient for the full lytic activity of vBcl-2, and that the N terminus of vBcl-2 likely coordinates multiple partially separable interactions, including those linked to tegument assembly [25], mitochondrial dynamics [26], and innate immune evasion [26].

The re-localization of KLHL9 to vBcl-2-positive mitochondrial regions is also conceptually important. vBcl-2 has already been implicated in mitochondrial remodeling through the NM23-H2-DRP1 axis and in suppression of MAVS-dependent interferon signaling [26]; the present findings therefore raise the possibility that recruitment of KLHL9 to mitochondria may represent an additional layer of this mitochondrial regulatory program. One attractive model is that vBcl-2 functions as a spatial organizer that redirects KLHL9, and potentially KLHL9/KLHL13-containing CRL3 complexes, toward mitochondrial or mitochondria-associated substrates that become important during lytic replication. Such substrates could be cellular proteins that restrict viral gene expression, organelle remodeling, innate immune signaling, or virion assembly. Alternatively, they could comprise viral proteins whose localization or activity must be regulated during infection. Consistent with this notion, we did not detect enhanced ubiquitination of vBcl-2 in the presence of KLHL9, even after proteasome inhibition with bortezomib or MG132. This negative result argues against a simple model in which vBcl-2 is directly recruited by KLHL9 as a degradative substrate, but rather suggests its ability to retarget host ubiquitin ligase activity to a subcellular compartment that is central to the lytic program. As BTB-Kelch adaptors can promote mono-or polyubiquitination, and as CUL3-based ligases can regulate substrate localization or function without necessarily inducing proteasomal degradation [45–47], KLHL9 may alter the activity, trafficking, or post-translational modification of another cellular or viral factor important for KSHV reactivation. This possibility is especially relevant in light of the known role of KLHL9/KLHL13 in Aurora B re-localization [35,36] and evidence that these adaptors can influence autophagy [38], mitochondrial quality control and antibacterial host response [39]. Identifying KLHL9-dependent ubiquitination events during lytic reactivation will be important for defining the mechanism of vBcl-2-mediated mitochondrial reprogramming and its overall contribution to successful viral replication.

The present study also helps reconcile several earlier observations concerning vBcl-2. Prior work showed that the canonical anti-apoptotic and anti-autophagic functions of vBcl-2 are dispensable for rescue of lytic replication, while distinct residues in the N terminus are required for productive infection [23]. Other studies linked this N terminal region to ORF55 binding [25], nuclear trafficking [24], and NM23-H2/DRP1-dependent mitochondrial fission [26]. Our findings suggest that KLHL9 should be considered part of this broader network of lytic vBcl-2 interactions. Rather than performing a single dedicated function, vBcl-2 may act as a multifunctional hub that assembles distinct protein complexes in different subcellular compartments and at different stages of the lytic cycle. Of note, CRLs are responsible for up to 20% of all ubiquitinated substrates within cells, and many viruses hijack or co-opt host CRLs by expressing proteins that physically interact with subunits of one or more CRL family members [48–51]. Furthermore, depletion of selected KLHL proteins, including KLHL13, decreases influenza A virus infection [52].

In summary, our results identify KLHL9 as a vBcl-2-associated host factor that supports efficient KSHV lytic replication. The data are consistent with a model in which vBcl-2 recruits KLHL9 to mitochondria, which likely to functionally engage a KLHL9-associated CRL3 complex to promote efficient lytic gene expression and virus production. Whether vBcl-2 only functions to recruit KLHL9 to the mitochondria or takes part in the CRL3 complex remains to be determined. More broadly, these findings strengthen the emerging view that gammaherpesvirus Bcl-2 homologs are not simply apoptosis and autophagy regulators, but multifunctional proteins that co-opt host trafficking, ubiquitin-signaling, and organelle-remodeling pathways to drive productive infection.

## Materials and Methods

### Cells

HEK-293T, SLK, and iSLK cells (kindly provided by Don Ganem and Rolf Renne) [53] were maintained in Dulbecco’s modified Eagle’s medium (DMEM) supplemented with 10% fetal bovine serum (FBS) (Gibco, Thermo Fisher Scientific), 50 IU/ml penicillin, and 50 µg/ml streptomycin (Biological Industries, Kibbutz Beit Haemek, Israel). 250 µg/ml of G418 and 1 µg/ml of puromycin (Formedium Ltd, England) were added to the iSLK cell growth medium to preserve the Tet-On transactivator and doxycycline-inducible RTA expression cassette, respectively. KSHV-infected HEK-293T and iSLK cells were maintained in the presence of 200 and 600 µg/mg hygromycin B (A. G. Scientific Incorporated), respectively. Lytic reactivation was induced in infected HEK-293T cells with 20 ng/ml 12-*O*-tetradecanoylphorbol-13-acetate (TPA) and 1 mM sodium butyrate (Sigma), and in infected iSLK cells with 1 µg/ml doxycycline and 1 mM sodium butyrate (Sigma). Infectious virus quantification was performed on clarified and filtered supernatants from induced producer cells to infect naive SLK cells followed by flow-cytometric measurement of GFP-positive cells at 48 h post infection, as previously described [54].

### Construction and reconstitution of recombinant BAC16-HA-vBcl-2 virus and quantification of infectious progeny

BAC16 containing a recombinant full-length KSHV genome carrying a green fluorescent protein (GFP) cassette under the control of the elongation factor 1α (EF-1α) promoter, has been described previously [42]. BAC16-HA-vBcl-2 was generated in *Escherichia coli* GS1783 using a two-step Red recombination strategy, as described previously [22,41]. An insertion cassette was amplified from pEGFP-N1 (Clontech) containing a kanamycin resistance (Kanr) cassette using primers designed to introduce an N-terminal HA tag at the ORF16/vBcl-2 initiation codon. The primer set used was 5′-ACACACCCTTGGTGCTGTGCGCGTGCTATGTGCCCTGGTGACCGTCCACA**ATG**<u>GC</u> <u>TTACCCATACGATGTTCCAGATTACGCT</u>GACGAGGACGTtagggataacagggtaatAGGTGGCACTTTTCGGGG-3′(the first 50 nucleotides [nt] correspond to nt 29,911 to 29,961 of BAC16, and are located upstream of ORF16, followed by an initiation codon (boldface) and 30 nt of the tag HA (underlined), 11 nt corresponding to nt 29,964-29,975 within ORF16, the I-SceI restriction site (lowercase letters) and the kanamycin resistance gene sequence) and 5′-CATGAATATCCCTTCAATGGCCAACACCTCTCCAGGCAAAACGTCCTCGTC <u>AGCGTAATCTGGAACATCGTATGGGTAAGC</u>**CAT**TTTATTGCCGTCATAGCGCGG-3′ (the first 51 nt correspond to nt 29,964-30,015 within *ORF16*, followed by 30 nt of the tag HA (underlined) and 21 nt corresponding to the kanamycin resistance gene sequence). Integration of the Kanr/I-SceI cassette was verified by PCR, followed by restriction enzyme analysis of purified BAC16 DNA. Diagnostic PCR was performed using primers located upstream of ORF16 (nt 29,941 to 29,960; 5′TGCCCTGGTGACCGTCCACA-3′) and within the ORF16 coding sequence (nt 30,423 to 30,408; 5′-TGTCATTCTCCGTCC-3′), yielding the expected 482 bp product. The integrated cassette was then excised upon treatment with 1% L-arabinose, enabling a second recombination event. BAC DNA was purified using a Large-Construct Kit (Qiagen) according to the manufacturer’s instructions. Parental and newly generated BAC DNAs were analyzed by BamHI digestion and run on a 0.4% agarose gel. The inserted DNA fragment was sequenced to confirm appropriate insertion of the tag.

BAC16 transfection and virus reconstitution were performed as previously described [22]. Briefly, BAC DNA was transfected into HEK-293T cells, followed by selection of cells harboring the viral genome, and co-cultivation of induced HEK-293T cells with iSLK. To prepare virions, supernatant from induced cells was collected and cleared of cells and debris by centrifugation (700 x *g* for 10 min at 4°C) and filtration (0.45-µm-pore-size cellulose acetate filters; Corning). Quantification of infectious virus production was performed as previously described [22].

### Plasmids and mutagenesis

The vBcl-2-coding sequence was cloned into a pcDNA vector containing an N-terminal hemagglutinin (HA) tag. Mutations in vBcl-2 gene were generated using Q5^®^ High-Fidelity DNA Polymerase (New England Biolabs). pcDNA3.1-KLHL9-Flag was obtained from GeneScript, whereas mCherry-KLHL9 and the corresponding deletion mutants were cloned in pDSRed. Single-guide RNAs (sgRNAs) targeting KLHL9 for use in iSLK cells were cloned into lentiCRISPRv2-Blast (Addgene). Lentiviral vectors encoding mCherry reporter only, and mCherry reporter and KLHL9-Flag with synonymous mutations that render the transcript resistant to sgRNA-mediated targeting was generated using VectorBuilder, pLV-mCherry-CMV-KLHL9-mut. sgRNAs targeting KLHL9 and KLHL13 for use in HEK293T cells were cloned into lentiCRISPRV2-Puro. Control cells were transduced with an AAVS1-targeting sgRNA construct.

### Antibodies

Primary antibodies used for western blot, immunoprecipitation and immunofluorescence analyses include: mouse anti-HA (Biolegend, MMS101R), mouse anti-RTA (kindly obtained from Prof. K. Ueda) [55], mouse anti-ORF45 (Santa Cruz, sc-53883), mouse anti-ORF K8 (Santa Cruz, sc-57889), mouse anti-ORF65 (Kindly obtained from Prof. S.J. Gao) [56] mouse anti-GFP (Santa Cruz, sc-9996), rabbit anti-mCherry (Abcam, 1C51), mouse anti-Flag (Sigma, F1804), mouse anti-Tubulin (DSHB, E7-s), and mouse anti-KLHL9/13 (Santa Cruz, sc-166486). Peroxidase-conjugated AffiniPure goat anti-mouse (Jackson-ImmunoResearch Laboratories, 115-035-062) was used as the secondary antibody for western blotting. For immunofluorescence analysis, Alexa Fluor 488 AffiniPure goat anti-mouse IgG (715-545-150), and Cy3 AffiniPure goat anti-rabbit IgG (111-165-045) were used as the secondary antibodies (Jackson-ImmunoResearch Laboratories).

### Immunoprecipitation, immunoblotting and proteomic analysis

For immunoprecipitation experiments, cells were lysed in RIPA buffer containing 20 mM Tris-HCl (pH 7.5), 150 mM NaCl, 1% NP40, 0.5% Deoxycholic acid, 2 mM Sodium vanadate, 1 mM Na_2_EDTA pH 8.0 and Complete Protease Inhibitor Cocktail (Roche). For each sample, 50 μg of clarified total protein extract was retained as input, and analyzed by SDS-PAGE and immunoblotting. For immunoprecipitation, approximately 500 μg of total protein extract was incubated overnight with the indicated antibody. Immune complexes were then captured by incubation with protein A/G Plus agarose beads (Santa Cruz Biotechnology). The beads were washed once with RIPA buffer and twice with PBS, and bound proteins were eluted by heating in 2× SDS sample buffer. Input and immunoprecipitated samples were resolved by SDS-PAGE and analyzed by immunoblotting with the indicated antibodies.

For proteomic identification of vBcl-2-associated proteins, immunoprecipitated samples were prepared as described above and analyzed by LC-MS/MS analysis (Smoler Proteomics Center, Technion). Samples were digested with trypsin, and analyzed on a Q Exactive mass spectrometer (Thermo Fisher Scientific). Peptide and protein identification was performed using Proteome Discoverer version 1.4 with two search algorithms, SEQUEST (Thermo Fisher Scientific) and Mascot (Matrix Science). Searches were performed against combined human and KSHV proteomes from the UniProt database, together with a decoy database to estimate the false discovery rate. Identified peptides were filtered to retain high-confidence, selecting top-ranked matches with appropriate mass accuracy and a minimum of two peptides per protein. High-confidence peptide identifications were defined as those passing a 1% false discovery rate threshold. Semiquantitative analysis was performed based on peptide peak areas, with protein abundance calculated as the average peak area of the three most intense peptides identified for each protein.

### Ubiquitination assay

To assay ubiquitination, cells were transfected with the indicated expression vectors, including Myc-Ubiquitin. Cells were treated with either 1 µM Bortezomib (Sigma) or 10 µM MG132 (Cell Signaling Technology), 6 hours before harvest. Cells were lysed in protein lysis buffer (PLB; 10 mM Tris/Cl pH 7.5, 150 mM NaCl, 0.5 mM EDTA, 0.5% NP40) containing the deubiquitinase inhibitor PR619 (ChromoTek). Immunoprecipitation was performed as described above.

### Immunofluorescence microscopy

Cells seeded on glass coverslips were washed with PBS and fixed with 4% PFA at room temperature, or 100% methanol at −20°C. After fixation, cells were washed again with PBS and blocked and permeabilized in PBS containing 0.2% Triton X-100 and 1% Bovine serum albumin (BSA, Sigma). Cells were then incubated with primary antibodies for 1-3 hr at room temperature, or overnight at 4°C, followed by incubation with fluorescently conjugated secondary antibodies for 1 hr at room temperature. Primary antibodies included mouse anti-HA (Biolegend, MMS101R) and rabbit anti-mCherry (Abcam) followed by Alexa Fluor 488 goat anti-mouse, Cy3 goat anti-rabbit secondary antibodies (Jackson Immuno Research Laboratories), respectively. For mitochondrial staining, cells were incubated with MitoTracker Deep Red FM (Cell Signaling, #8778) according to the manufacturer’s instructions, followed by immunostaining, as described above. Nuclei were stained with 0.05 µg/ml Hoechst (Sigma) in PBS. Coverslips were mounted with antifade mounting solution containing 90% glycerol, 9% PBS and 1% n-propyl-gallate (Sigma). Images were acquired on a Leica confocal LIVE microscope. The fluorescence intensity profile of images was quantified using ImageJ.

### Generation of KLHL9-knockout iSLK and HEK293T cells by CRISPR/Cas9

KLHL9 knockout (KO) in BAC16-HA-vBcl-2-containing iSLK cells was generated using a lentiviral CRISPR/Cas9 approach with lentiCRISPRv2-Blast vector (Addgene). The single-guide RNA (sgRNA) sequence 5′-CACCGCACAGTTCGGTGGTATTGCA-3’ was used to successfully disrupt KLHL9. A corresponding control vector was used in parallel. Following lentiviral transduction, iSLK-BAC16-HA-vBcl-2 cells were selected with 15 µg/ml blasticidin to generate a stable KLHL9-knockout cell population. Disruption of the KLHL9 locus was confirmed by PCR amplification of the genomic region encompassing the sgRNA target site, followed by DNA sequencing. HEK293T cells deficient in KLHL9 and KLHL13 were generated using the CRISPR/Cas9 system. sgRNA sequences targeting KLHL9 or KLHL13 were cloned into the lentiCRISPR v2-Puro vector (Addgene). The sgRNA sequences used were as follows: sgAAVS1, 5′-GGGGCCACTAGGGACAGGAT-3′; sgKLHL9, 5′-GATCTCCATCACCTGGTACCA-3′; and sgKLHL13, 5′-GACAACTGTGTTGAAGTTGGA-3′. Cells expressing an sgRNA targeting the AAVS1 safe-harbor locus were used as controls. Following lentiviral transduction, cells were selected in growth medium containing 1 µg/ml puromycin. Single-cell-derived clones were isolated and screened by genomic sequencing. Editing efficiency and indel patterns were analyzed using Synthego ICE analysis.

### Complementation of KLHL9-knockout iSLK cells

For complementation, iSLK-BAC16-HA-vBcl-2-KLHL9-KO cells were transduced with the lentiviral expression vector, pLV-mCherry-CMV-KLHL9-mut (VectorBuilder), which co-expresses mCherry and a KLHL9 form carrying synonymous substitutions that render it resistant to CRISPR/Cas9 targeting. The complemented KLHL9 protein was tagged at its C terminus with FLAG. As a control, cells were transduced with the corresponding empty vector, pLV-CMV-mCherry, which expresses mCherry alone. Approximately two weeks after transduction, mCherry-positive cells were enriched by fluorescence-activated cell sorting (FACS) using a FACSAria III cell sorter to obtain a homogeneous population.

### Structural prediction and analysis of the vBcl-2-KLHL9 protein complex

To predict the protein complex of KSHV vBcl-2 and KLHL9, we used AlphaFold-Multimer with default parameters [57]. The highest-scoring protein complex had a prediction score of 0.788, which is considered to be high (and is based on the summation of 0.8×ipTM+0.2×pTM). Using this structural model, we employed the contacts of structural units (CSU) method [58], which identifies protein residues that come into spatial contact with each other, allowing identification of interface residues. To visualize the results, images of the structure were created using PyMol (https://www.pymol.org/).

## Acknowledgements

We gratefully acknowledge Don Ganem, Rolf Renne, Jae. U. Jung, Shou Jiang Gao and Keiji Ueda for reagents.

This work was supported by a grant from the Israel Science Foundation (ISF) (grant no. 1932/23) to R.S., and by the Ephraim Rapaport Chair in Clinical and Medical Microbiology (R.S).

---

**<u>S1 Table</u>. Identification of host and viral proteins associated with HA-vBcl-2 by LC-MS/MS.** iSLK cells harboring wild-type BAC16 or BAC16-HA-vBcl-2 were induced into the lytic cycle with 1 µg/ml doxycycline and 1 mM sodium butyrate. Cells were harvested 48 h following induction, and protein lysates were subjected to immunoprecipitation with anti-HA antibody followed by LC-MS/MS analysis. Peptides were identified using Proteome Discoverer against the human and KSHV proteomes, and proteins were semi-quantified based on the average peak area of the three most intense peptides. Proteins enriched by at least 5-fold in BAC16-HA-vBcl-2 samples relative to wild-type BAC16 controls were considered candidate interacting partners.

**<u>S2 Table</u>**. **List of interface contacts between vBcl-2 and KLHL9.** Interacting residues between the two proteins was determined using the contacts of structural units (CSU) approach, as detailed in Methods section.

## References

1. Chang Y, Cesarman E, Pessin MS, Lee F, Culpepper J, Knowles DM, et al. Identification of Herpesvirus-Like DNA Sequences in AIDS-Sssociated Kaposi’s Sarcoma. Science. 1994;266: 1865–1869. doi:10.1126/science.7997879

2. Mariggiò G, Koch S, Schulz TF. Kaposi sarcoma herpesvirus pathogenesis. Phil Trans R Soc B. 2017;372: 20160275. doi:10.1098/rstb.2016.0275

3. Calabrò ML, Sarid R. Human Herpesvirus 8 and Lymphoproliferative Disorders. Mediterr J Hematol Infect Dis. 2018;10: e2018061. doi:10.4084/MJHID.2018.061

4. Cesarman E, Damania B, Krown SE, Martin J, Bower M, Whitby D. Kaposi sarcoma. Nat Rev Dis Primers. 2019;5: 9. doi:10.1038/s41572-019-0060-9

5. Jary A, Veyri M, Gothland A, Leducq V, Calvez V, Marcelin A-G. Kaposi’s Sarcoma-Associated Herpesvirus, the Etiological Agent of All Epidemiological Forms of Kaposi’s Sarcoma. Cancers. 2021;13: 6208. doi:10.3390/cancers13246208

6. Oksenhendler E, Meignin V. HHV-8 associated lymphoma. Current Opinion in Oncology. 2022;34: 432–438. doi:10.1097/CCO.0000000000000884

7. Damania B, Dittmer DP. Today’s Kaposi sarcoma is not the same as it was 40 years ago, or is it? J Med Virol. 2023;95: e28773. doi:10.1002/jmv.28773

8. Purushothaman P, Uppal T, Verma SC. Molecular biology of KSHV lytic reactivation. Viruses. 2015;7: 116–153. doi:10.3390/v7010116

9. Yan L, Majerciak V, Zheng Z-M, Lan K. Towards Better Understanding of KSHV Life Cycle: from Transcription and Posttranscriptional Regulations to Pathogenesis. Virol Sin. 2019;34: 135–161. doi:10.1007/s12250-019-00114-3

10. Murdock SJ, Bersonda JR, Forrest JC, Manzano M. Molecular Mechanisms of KSHV Latency Establishment and Maintenance. Curr Clin Micro Rpt. 2024;11: 220–230. doi:10.1007/s40588-024-00232-x

11. Losay VA, Damania B. Unraveling the Kaposi Sarcoma-Associated Herpesvirus (KSHV) Lifecycle: An Overview of Latency, Lytic Replication, and KSHV-Associated Diseases. Viruses. 2025;17: 177. doi:10.3390/v17020177

12. Cheng EH-Y, Nicholas J, Bellows DS, Hayward GS, Guo H-G, Reitz MS, et al. A Bcl-2 homolog encoded by Kaposi sarcoma-associated virus, human herpesvirus 8, inhibits apoptosis but does not heterodimerize with Bax or Bak. Proc Natl Acad Sci USA. 1997;94: 690–694. doi:10.1073/pnas.94.2.690

13. Sarid R, Sato T, Bohenzky RA, Russo JJ, Chang Y. Kaposi’s sarcoma-associated herpesvirus encodes a functional Bcl-2 homologue. Nat Med. 1997;3: 293–298. doi:10.1038/nm0397-293

14. Huang Q, Petros AM, Virgin HW, Fesik SW, Olejniczak ET. Solution structure of a Bcl-2 homolog from Kaposi sarcoma virus. Proc Natl Acad Sci USA. 2002;99: 3428–3433. doi:10.1073/pnas.062525799

15. Loh J, Huang Q, Petros AM, Nettesheim D, Van Dyk LF, Labrada L, et al. A Surface Groove Essential for Viral Bcl-2 Function During Chronic Infection In Vivo. McFadden G, editor. PLoS Pathog. 2005;1: e10. doi:10.1371/journal.ppat.0010010

16. Flanagan AM, Letai A. BH3 domains define selective inhibitory interactions with BHRF-1 and KSHV BCL-2. Cell Death Differ. 2008;15: 580–588. doi:10.1038/sj.cdd.4402292

17. Suraweera CD, Hinds MG, Kvansakul M. Structural Insight into KsBcl-2 Mediated Apoptosis Inhibition by Kaposi Sarcoma Associated Herpes Virus. Viruses. 2022;14: 738. doi:10.3390/v14040738

18. Bellows DS, Chau BN, Lee P, Lazebnik Y, Burns WH, Hardwick JM. Antiapoptotic Herpesvirus Bcl-2 Homologs Escape Caspase-Mediated Conversion to Proapoptotic Proteins. J Virol. 2000;74: 5024–5031. doi:10.1128/JVI.74.11.5024-5031.2000

19. Pattingre S, Tassa A, Qu X, Garuti R, Liang XH, Mizushima N, et al. Bcl-2 Antiapoptotic Proteins Inhibit Beclin 1-Dependent Autophagy. Cell. 2005;122: 927–939. doi:10.1016/j.cell.2005.07.002

20. Wei Y, Pattingre S, Sinha S, Bassik M, Levine B. JNK1-Mediated Phosphorylation of Bcl-2 Regulates Starvation-Induced Autophagy. Molecular Cell. 2008;30: 678–688. doi:10.1016/j.molcel.2008.06.001

21. Liang C, E. X, Jung JU. Downregulation of autophagy by herpesvirus Bcl-2 homologs. Autophagy. 2008;4: 268–272. doi:10.4161/auto.5210

22. Gelgor A, Kalt I, Bergson S, Brulois KF, Jung JU, Sarid R. Viral Bcl-2 Encoded by the Kaposi’s Sarcoma-Associated Herpesvirus Is Vital for Virus Reactivation. Longnecker RM, editor. J Virol. 2015;89: 5298–5307. doi:10.1128/JVI.00098-15

23. Liang Q, Chang B, Lee P, Brulois KF, Ge J, Shi M, et al. Identification of the Essential Role of Viral Bcl-2 for Kaposi’s Sarcoma-Associated Herpesvirus Lytic Replication. Longnecker RM, editor. J Virol. 2015;89: 5308–5317. doi:10.1128/JVI.00102-15

24. Gallo A, Lampe M, Günther T, Brune W. The Viral Bcl-2 Homologs of Kaposi’s Sarcoma-Associated Herpesvirus and Rhesus Rhadinovirus Share an Essential Role for Viral Replication. Frueh K, editor. J Virol. 2017;91: e01875–16. doi:10.1128/JVI.01875-16

25. Liang Q, Wei D, Chung B, Brulois KF, Guo C, Dong S, et al. Novel Role of vBcl2 in the Virion Assembly of Kaposi’s Sarcoma-Associated Herpesvirus. Sandri-Goldin RM, editor. J Virol. 2018;92: e00914–17. doi:10.1128/JVI.00914-17

26. Zhu Q, McElroy R, Machhar JS, Cassel J, Zheng Z, Mansoori B, et al. Kaposi’s sarcoma-associated herpesvirus induces mitochondrial fission to evade host immune responses and promote viral production. Nat Microbiol. 2025;10: 1501–1520. doi:10.1038/s41564-025-02018-3

27. Sarikas A, Hartmann T, Pan Z-Q. The cullin protein family. Genome Biol. 2011;12: 220. doi:10.1186/gb-2011-12-4-220

28. Duda DM, Scott DC, Calabrese MF, Zimmerman ES, Zheng N, Schulman BA. Structural regulation of cullin-RING ubiquitin ligase complexes. Current Opinion in Structural Biology. 2011;21: 257–264. doi:10.1016/j.sbi.2011.01.003

29. Chen H-Y, Chen R-H. Cullin 3 Ubiquitin Ligases in Cancer Biology: Functions and Therapeutic Implications. Front Oncol. 2016;6. doi:10.3389/fonc.2016.00113

30. Jerabkova K, Sumara I. Cullin 3, a cellular scripter of the non-proteolytic ubiquitin code. Seminars in Cell & Developmental Biology. 2019;93: 100–110. doi:10.1016/j.semcdb.2018.12.007

31. Genschik P, Sumara I, Lechner E. The emerging family of CULLIN3-RING ubiquitin ligases (CRL3s): cellular functions and disease implications. EMBO J. 2013;32: 2307–2320. doi:10.1038/emboj.2013.173

32. Dhanoa BS, Cogliati T, Satish AG, Bruford EA, Friedman JS. Update on the Kelch-like (KLHL) gene family. Hum Genomics. 2013;7: 13. doi:10.1186/1479-7364-7-13

33. Cirak S, Von Deimling F, Sachdev S, Errington WJ, Herrmann R, Bönnemann C, et al. Kelch-like homologue 9 mutation is associated with an early onset autosomal dominant distal myopathy. Brain. 2010;133: 2123–2135. doi:10.1093/brain/awq108

34. Izumi R, Fukasaka I, Matsumura T, Nakamura N, Takahashi T, Suzuki N, et al. KLHL9-linked distal myopathy: a second family suggesting broad phenotypic variability. Neuromuscular Disorders. 2025;55: 105450. doi:10.1016/j.nmd.2025.105450

35. Sumara I, Quadroni M, Frei C, Olma MH, Sumara G, Ricci R, et al. A Cul3-Based E3 Ligase Removes Aurora B from Mitotic Chromosomes, Regulating Mitotic Progression and Completion of Cytokinesis in Human Cells. Developmental Cell. 2007;12: 887–900. doi:10.1016/j.devcel.2007.03.019

36. Krupina K, Kleiss C, Metzger T, Fournane S, Schmucker S, Hofmann K, et al. Ubiquitin Receptor Protein UBASH3B Drives Aurora B Recruitment to Mitotic Microtubules. Developmental Cell. 2016;36: 63–78. doi:10.1016/j.devcel.2015.12.017

37. Chen JC, Alvarez MJ, Talos F, Dhruv H, Rieckhof GE, Iyer A, et al. Identification of Causal Genetic Drivers of Human Disease through Systems-Level Analysis of Regulatory Networks. Cell. 2014;159: 402–414. doi:10.1016/j.cell.2014.09.021

38. Begun J, Lassen KG, Jijon HB, Baxt LA, Goel G, Heath RJ, et al. Integrated Genomics of Crohn’s Disease Risk Variant Identifies a Role for CLEC12A in Antibacterial Autophagy. Cell Reports. 2015;11: 1905–1918. doi:10.1016/j.celrep.2015.05.045

39. Nan D, Rao C, Tang Z, Yang W, Wu P, Chen J, et al. Burkholderia pseudomallei BipD modulates host mitophagy to evade killing. Nat Commun. 2024;15: 4740. doi:10.1038/s41467-024-48824-x

40. Frendo-Cumbo S, Jaldin-Fincati JR, Coyaud E, Laurent EMN, Townsend LK, Tan JMJ, et al. Deficiency of the autophagy gene ATG16L1 induces insulin resistance through KLHL9/KLHL13/CUL3-mediated IRS1 degradation. Journal of Biological Chemistry. 2019;294: 16172–16185. doi:10.1074/jbc.RA119.009110

41. Karstentischer B, Von Einem J, Kaufer B, Osterrieder N. Two-Step Red-Mediated Recombination for versatile High-Efficiency Markerless DNA Manipulation in *Escherichia Coli*. BioTechniques. 2006;40: 191–197. doi:10.2144/000112096

42. Brulois KF, Chang H, Lee AS-Y, Ensser A, Wong L-Y, Toth Z, et al. Construction and Manipulation of a New Kaposi’s Sarcoma-Associated Herpesvirus Bacterial Artificial Chromosome Clone. J Virol. 2012;86: 9708–9720. doi:10.1128/JVI.01019-12

43. Davis ZH, Verschueren E, Jang GM, Kleffman K, Johnson JR, Park J, et al. Global Mapping of Herpesvirus-Host Protein Complexes Reveals a Transcription Strategy for Late Genes. Molecular Cell. 2015;57: 349–360. doi:10.1016/j.molcel.2014.11.026

44. Yao Y, Hong S, Yoshida S, Swaroop V, Curtin B, Inoki K. The Cullin3-Rbx1-KLHL9 E3 ubiquitin ligase complex ubiquitinates Rheb and supports amino acid-induced mTORC1 activation. Cell Reports. 2025;44: 115101. doi:10.1016/j.celrep.2024.115101

45. Yuan W-C, Lee Y-R, Lin S-Y, Chang L-Y, Tan YP, Hung C-C, et al. K33-Linked Polyubiquitination of Coronin 7 by Cul3-KLHL20 Ubiquitin E3 Ligase Regulates Protein Trafficking. Molecular Cell. 2014;54: 586–600. doi:10.1016/j.molcel.2014.03.035

46. Werner A, Iwasaki S, McGourty CA, Medina-Ruiz S, Teerikorpi N, Fedrigo I, et al. Cell-fate determination by ubiquitin-dependent regulation of translation. Nature. 2015;525: 523–527. doi:10.1038/nature14978

47. Ramirez-Martinez A, Cenik BK, Bezprozvannaya S, Chen B, Bassel-Duby R, Liu N, et al. KLHL41 stabilizes skeletal muscle sarcomeres by nonproteolytic ubiquitination. eLife. 2017;6: e26439. doi:10.7554/eLife.26439

48. Mahon C, Krogan N, Craik C, Pick E. Cullin E3 Ligases and Their Rewiring by Viral Factors. Biomolecules. 2014;4: 897–930. doi:10.3390/biom4040897

49. Li S, Li R, Ahmad I, Liu X, Johnson SF, Sun L, et al. Cul3-KLHL20 E3 ubiquitin ligase plays a key role in the arms race between HIV-1 Nef and host SERINC5 restriction. Nat Commun. 2022;13: 2242. doi:10.1038/s41467-022-30026-y

50. Ding S, Mooney N, Li B, Kelly MR, Feng N, Loktev AV, et al. Comparative Proteomics Reveals Strain-Specific β-TrCP Degradation via Rotavirus NSP1 Hijacking a Host Cullin-3-Rbx1 Complex. Coyne CB, editor. PLoS Pathog. 2016;12: e1005929. doi:10.1371/journal.ppat.1005929

51. Lutz LM, Pace CR, Arnold MM. Rotavirus NSP1 Associates with Components of the Cullin RING Ligase Family of E3 Ubiquitin Ligases. López S, editor. J Virol. 2016;90: 6036–6048. doi:10.1128/JVI.00704-16

52. Gschweitl M, Ulbricht A, Barnes CA, Enchev RI, Stoffel-Studer I, Meyer-Schaller N, et al. A SPOPL/Cullin-3 ubiquitin ligase complex regulates endocytic trafficking by targeting EPS15 at endosomes. eLife. 2016;5: e13841. doi:10.7554/eLife.13841

53. Myoung J, Ganem D. Generation of a doxycycline-inducible KSHV producer cell line of endothelial origin: Maintenance of tight latency with efficient reactivation upon induction. Journal of Virological Methods. 2011;174: 12–21. doi:10.1016/j.jviromet.2011.03.012

54. Orbaum-Harel O, Sloutskin A, Kalt I, Sarid R. KSHV ORF20 Promotes Coordinated Lytic Reactivation for Increased Infectious Particle Production. Viruses. 2024;16: 1418. doi:10.3390/v16091418

55. Ueda K, Ishikawa K, Nishimura K, Sakakibara S, Do E, Yamanishi K. Kaposi’s Sarcoma-Associated Herpesvirus (Human Herpesvirus 8) Replication and Transcription Factor Activates the K9 (vIRF) Gene through Two Distinct *cis* Elements by a Non-DNA-Binding Mechanism. J Virol. 2002;76: 12044–12054. doi:10.1128/JVI.76.23.12044-12054.2002

56. Ye F, Zhou F, Bedolla RG, Jones T, Lei X, Kang T, et al. Reactive Oxygen Species Hydrogen Peroxide Mediates Kaposi’s Sarcoma-Associated Herpesvirus Reactivation from Latency. Früh K, editor. PLoS Pathog. 2011;7: e1002054. doi:10.1371/journal.ppat.1002054

57. Evans R, O’Neill M, Pritzel A, Antropova N, Senior A, Green T, et al. Protein complex prediction with AlphaFold-Multimer. Bioinformatics; 2021. doi:10.1101/2021.10.04.463034

58. Sobolev V, Sorokine A, Prilusky J, Abola EE, Edelman M. Automated analysis of interatomic contacts in proteins. Bioinformatics. 1999;15: 327–332. doi:10.1093/bioinformatics/15.4.327

